# High-Throughput Characterization of Protein-Protein Interactions by Reprogramming Yeast Mating

**DOI:** 10.1101/122143

**Authors:** David Younger, Stephanie Berger, David Baker, Eric Klavins

## Abstract

High-throughput methods for screening protein-protein interactions enable the rapid characterization of engineered binding proteins and interaction networks. While existing approaches are powerful, none allow quantitative library-on-library characterization of protein interactions in a modifiable extracellular environment. Here, we show that sexual agglutination of *S. cerevisiae* can be reprogrammed to link interaction strength with mating efficiency using synthetic agglutination (SynAg). Validation of SynAg with 89 previously characterized interactions shows a log-linear relationship between mating efficiency and protein binding strength for interactions with K_D_’s ranging from below 500 pM to above 300 μM. Using induced chromosomal translocation to pair barcodes representing binding proteins, thousands of distinct interactions can be screened in a single pot. We demonstrate the ability to characterize protein interaction networks in a modifiable environment by introducing a soluble peptide that selectively disrupts a subset of interactions in a representative network by up to 800-fold. SynAg enables the high-throughput, quantitative characterization of protein-protein interaction networks in a fully-defined extracellular environment at a library-on-library scale.

**Significance Statement:** *De novo* engineering of protein binders often requires experimental screening to select functional variants from a design library. We have achieved high-throughput, quantitative characterization of protein-protein binding interactions without requiring purified recombinant proteins, by linking interaction strength with yeast mating. Using a next-generation sequencing output, we have characterized protein networks consisting of thousands of pairwise interactions in a single tube and have demonstrated the effect of changing the binding environment. This approach addresses an existing bottleneck in protein binder design by enabling the high-throughput and quantitative characterization of binding strength between designed protein libraries and multiple target proteins in a fully defined environment.

## Introduction

Many powerful methods have been developed for the high-throughput screening of protein-protein interactions. Yeast two-hybrid^1^ can be used to intracellularly screen pairwise interactions and has been extended to the screening of large protein interaction networks using next-generation sequencing^2,3,4^. However, intracellular assays are limited by a lack of control over the binding environment and suffer from frequent false-positives and false-negatives^5,6^ Phage^7^ and yeast^8^ display have enabled the high-throughput binding characterization of large protein libraries, but can only screen binding against a limited number of targets simultaneously due to the spectral resolution of existing fluorescent reporters^9^. Both approaches also require the expression and purification of recombinant target proteins, which is limiting if the target is unstable or expresses poorly. Single molecular interaction sequencing^10^ can characterize protein interaction networks in a cell-free environment, but requires purified proteins and a dedicated flow cell for the analysis of each network and condition. While each approach expands protein interaction screening capabilities, none allows for cell-based, quantitative, library-on-library interaction characterization in an extracellular environment that can be modified as desired.

Yeast mating in an aerated liquid culture depends critically on an intercellular protein-protein interaction that drives agglutination between MATa and MATα haploid cells^11,12^. The MATa sexual agglutinin subunit, Aga2, and the MATα cognate, Sag1, interact with a K_D_ of 2 to 5 nM and initiate the irreversible binding of two haploid cells and subsequent cellular fusion to create a single diploid cell^13^. Cellular agglutination and mating is highly efficient, occurs in a matter of hours, and each mating event forms a stable and propagating diploid strain^14^.

Here, we reprogram yeast mating by knocking out Sag1 and replacing wild-type agglutination with engineered or natural binding pairs that act as synthetic agglutination (SynAg) proteins. SynAg proteins are expressed on the cell surface as fusions to Aga2 using yeast surface display, which has been used to functionally display proteins with diverse structural properties, such as single-chain antibody fragments^15^, green fluorescent protein^16^, and the human epidermal growth factor receptor ectodomain^17^. Thus, we expect that SynAg will be broadly compatible with soluble proteins of interest. We first show that interaction strength between one MATa SynAg protein and one MATα SynAg protein can be quantitatively assessed by co-culturing the two haploid strains and measuring their mating efficiency with a flow cytometry assay (Fig. 1a). We then extend the approach to one-pot, library-on-library characterization by barcoding SynAg gene cassettes, co-culturing many MATa and MATα strains, and using next-generation sequencing to count interaction frequencies for all possible MATa-MATα SynAg protein interactions (Fig. 1b). Finally, we demonstrate that the binding environment can be systematically manipulated to probe the influence of external variables on protein interaction networks.

**Figure 1.**
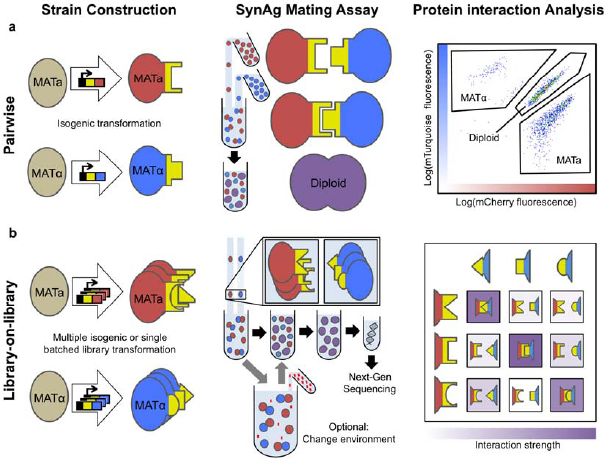
Yeast synthetic agglutination (SynAg) workflow. **(a)** Pairwise characterization of a protein interaction involves isogenic yeast transformations followed by a SynAg mating assay and flow cytometry, **(b)** Library-on-library protein interaction network characterization involves library construction followed by a SynAg mating assay, diploid DNA isolation, and next-generation sequencing. Changing the binding environment during the SynAg mating assay is optional.

## Results

### Reprogramming Sexual Agglutination

In order to characterize protein-protein interactions using yeast mating, we first genetically replaced native sexual agglutination with SynAg interactions. To validate our approach, we used members of the well-characterized BCL2 family^18^, which regulate apoptosis in human cells via binding interactions between pro-survival and pro-apoptotic family members, as SynAg proteins (Table S1). Six pro-survival BCL2 homologues were expressed on MATα cells. Binding peptides from seven pro-apoptotic BCL2 proteins were expressed on MATa cells. Previously, the pairwise interaction strength between the pro-survival homologues and pro-apoptotic peptides were semi-quantitatively characterized using a competition binding assay that showed over 10,000-fold differences in interaction strength^19^. Nine *de novo* binding proteins, also exhibiting a wide range in interaction strength for the pro-survival homologues, were expressed on MATα cells. Previously, the pairwise interaction strength between pro-survival homologues and engineered binders were quantitatively characterized using biolayer interferometry^20,21^.

Isogenic yeast strains were generated for the expression of each SynAg protein by transforming Sag1 deficient strains with a fragment containing a SynAg surface expression cassette and a mating type-specific fluorescent reporter. Pairs of SynAg-expressing haploid cells were co-cultured in non-selective liquid media for 17 hours to allow agglutination-dependent mating. Flow cytometry was performed to differentiate between mCherry expressing MATa haploids, mTurquoise expressing MATα haploids, and diploids that expressed both fluorescent markers (Fig. 1a). Diploid percent was used as a metric for mating efficiency to quantitatively characterize the interaction strength between a MATa SynAg protein and a MATα SynAg protein.

We found that complementary SynAg proteins expressed on the surface of yeast are necessary and sufficient to replace the function of Aga2 and Sag1. Wild-type *S. cerevisiae* haploid cells mated with an efficiency of 63.6% ± 3.1% in standard laboratory conditions and a knockout of Sag1 in the MATα haploid eliminated mating with wild-type MATa (Fig. 2a). In the Sag1 knockout, expression of an interacting SynAg protein pair recovered mating efficiency to 51.6% ± 7.9%, while expression of a non-interacting SynAg protein pair showed no observable recovery (Fig. 2b). SynAg-dependent recovery of mating occurred with a variety of natural and engineered proteins and peptides ranging from 26 to 206 amino acids, indicating a large engineerable space for synthetic agglutination.

**Figure 2.**
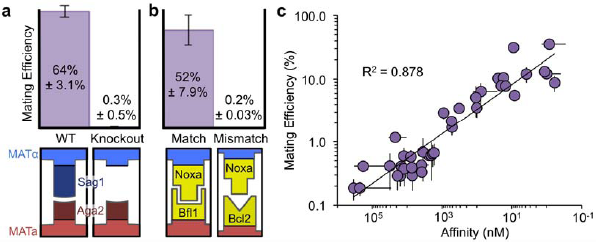
SynAg for characterizing pairwise protein interactions. **(a)** The mating efficiency of wild-type *S. cerevisiae* haploid cells is eliminated with the knockout of the MATα sexual agglutinin protein, Sag1. **(b)** Complementary binding proteins expressed on the surface of yeast haploid cells recover mating efficiency, **(c)** Mating efficiencies for SynAg protein pairs with a K_D_ from below 500 pM to above 300 μM show a log-linear relationship with affinity, measured with biolayer interferometry (SD, n=3)^21^.

### Pairwise Characterization of Protein Interactions

Mating efficiency and affinity, measured with biolayer interferometry, are related log-linearly (R^2^=0.878) for protein interactions across over five orders of magnitude of K_D_ (Fig. 2c). We tested proteins with binding affinities ranging from below 500 pM to above 300 μM, which gave mating efficiencies up to 35.4% and down to below 0.2%. The strong log-linear relationship between mating efficiency and affinity over multiple orders of magnitude contradicted our expectation of avidity as the main driving force for yeast agglutination^13^. We expected that upon the formation of a single interaction between cells, newly localized protein pairs would rapidly bind, making off-rate largely irrelevant. However, both on-and off-rate showed a correlation with mating efficiency, and neither provided as good a fit as K_D_ (Fig. S2a,b).

SynAg proteins must be adequately expressed on the cell surface in order to accurately correlate mating efficiency and affinity. To test the effect of surface expression strength on mating efficiency, we constructed an inducible SynAg expression cassette with a synthetic transcription factor^22^ that could be tuned to different levels by changing the inducer concentration (Fig. S2c). Surface expression strength was measured by labeling with FITC-conjugated anti-myc^8^and measuring fluorescence with flow-cytometry. While mating efficiency was highly dependent on surface expression strength at low levels of expression, the effect saturated above 4,000 au (Fig. S2d). Of all SynAg proteins tested, one BCL2 homologue showed surface expression strength below 4,000 au and subsequently minimal recovery of mating efficiency regardless of its mating partner (Fig. S2e). An alternative semi-functional truncation improved surface expression and enabled affinity-dependent mating^23^ (Fig. S2f).

### Barcoding and recombination of interaction libraries

A chromosomal barcoding and recombination scheme was developed for one-pot library-on-library protein interaction network characterization, which uses a similar approach to previous work on plasmid-based recombination^3^. We began by constructing MATa and MATα parent strains, ySYNAGa and ySYNAGα, into which SynAg expression cassette libraries were transformed (Fig. S3). These strains contain complementary lysine and leucine auxotrophic markers for diploid selection and express CRE recombinase^24^ after mating when induced with (β-Estradiol^25^. For small libraries, SynAg cassettes were assembled with isothermal assembly^26^ in one of two standardized vectors, pSYNAGa or pSYNAGα, for integration into yeast (Fig. S4a,b). In addition to the surface expression cassette, each vector backbone contains a randomized SynAg protein specific barcode and a mating type specific lox recombination site and primer binding site. Sanger sequencing^27^ was used to match barcodes with their corresponding SynAg proteins. The construction of SynAg libraries is comparable in time and cost to the construction of yeast two-hybrid or yeast surface display libraries and identical methods can be used for DNA preparation and transformation.

Unidirectional CRE induced chromosomal translocation^28^ in diploid cells resulted in the combining of barcodes representing two interacting SynAg proteins onto the same chromosome (Fig. 3a). After recombination, interacting SynAg proteins were identified from a mixed culture using Illumina next-generation sequencing^29^. To test the approach, haploid cells containing SynAg cassettes were mated, induced with β-Estradiol, and lysed. The yeast lysate was used as a template for a PCR using mating type-specific primers. Amplification indicated that the primer binding sites had been combined onto one contiguous DNA strand, and hence that recombination had occurred. Sanger sequencing of the amplicon confirmed that recombination resulted in the expected chromosomal translocation.

**Figure 3.**
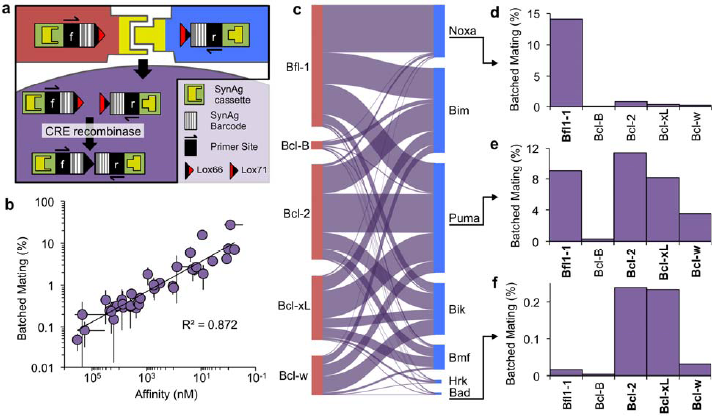
SynAg for one-pot library-on-library protein network characterization. **(a)** Each SynAg protein expression cassette is flanked by a barcode unique to a particular SynAg protein and a mating type specific primer binding site and lox recombination site. CRE recombinase expression in diploid cells combines MATa and MATα barcodes onto the same chromosome for next-generation sequencing, **(b)** A one-pot library-on-library mating assay gives a strong log-linear relationship between batched mating percent and affinity for protein interactions with a K_D_from below 500 pM to above 300 μM (SD, n=2). **(c)** All interactions between five pro-survival BCL2 homologues and seven pro-apoptotic peptides (BH3 domains of Noxa, Bim, Puma, etc.) were characterized with a library-on-library SynAg assay. The height of each purple bar represents the frequency of diploid formation from a given pair of SynAg expressing haploid strains. **(d,e,f)**Interaction profiles for **(d)** BH3 domain of Noxa (Noxa.BH3), **(e)** Puma.BH3, and **(f)** Bad.BH3 are shown in greater detail, with bolded BCL2 homologues indicating an affinity below 1 μM, according to a competition assay^19^.

### Library-on-library characterization of protein interactions

The frequency with which pairs of barcodes corresponding to interacting SynAg proteins appear in diploid lysate following a library-on-library mating was observed to be log-linear with biolayer interferometry affinity measurements (R^2^ = 0.872) (Fig. 3b). As before, we validated our approach with previously characterized protein interactions involving six pro-survival BCL2 homologues and nine *de novo* binding proteins. Here, all 15 SynAg strains were combined in a single mating. The batched mating percent for each interaction in the combinatorial matrix was calculated from next-generation sequencing counts, providing a relative strength for each protein interaction. We found that protein interactions spanning five orders of magnitude of K_D_ led to a more than 500-fold difference in batched mating percent. In addition to the *de novo* binding proteins, seven pro-apoptotic peptides with diverse binding profiles for the pro-survival homologues were added to a batched mating^30^ and the observed interactions were consistent with previous work^19^ (Fig. 3c,d,e,f).

A comparison of the pairwise and library-on-library SynAg methods showed a near perfect 1:1 agreement (Fig. S5). To compare the two approaches, pairwise mating efficiency was normalized so that the mating efficiency of all tested pairs summed to 100. A paired two-sided T-test of normalized pairwise mating percent and batched mating percent gave a p-value of 0.80, indicating no statistically significant difference between the two methods.

### Large protein-protein interaction library characterization

For the construction of large chromosomally integrated libraries, a “landing pad” approach^31^ was used to achieve high efficiency transformations from an integration requiring four-fragment homologous recombination (Fig. S4c). At the SynAg cassette integration locus, both ySYNAGa and ySYNAGα were transformed with an expression cassette consisting of Seel endonuclease driven by a galactose-inducible promoter and flanked by Seel cut sites. Galactose induction prior to transformation with a SynAg protein library resulted in DNA nicking at the site of integration, which dramatically improved transformation efficiency^32^. Next-generation sequencing of genomic DNA extracted from transformed yeast libraries was used to pair each SynAg protein variant to its distinct barcode and to count relative barcode frequencies in the naïve library prior to mating.

A single-pot batched mating was used to characterize 7,000 distinct protein-protein interactions. A partial site-saturation mutagenesis library of a *de novo* Bcl-xL binding protein, XCDP07^21^, consisting of 1,400 distinct variants was characterized for interactions with five pro¬survival BCL2 homologues. For each variant, interaction strength (the number of times a particular variant was observed to have mated with Bcl-xL divided by the number of times that variant was observed in the naïve library) and specificity (the percent of observed matings with Bcl-xL minus the percent of observed matings with the next highest homologue) were determined. As a proof of principal, interactions involving SynAg variants with premature stop codons were analyzed (Fig. 4a). Only 8 of 55 premature stop codons included in the library resulted in even a single mating and only 6 resulted in more than 2 matings. These six variants contained stop codons at residue 93 or later, which leaves the central binding helix intact. Two variants, with stop codons at residues 113 and 114, showed improved interaction strength and specificity for Bcl-xL. These early terminations resulted in the removal of the C-terminal myc tag from the 116-residue full-length protein, which may have negatively impacted binding. The same analysis was repeated with similar results for a site-saturation mutagenesis library of a Bcl-2 binder, 2CDP06 (Fig. S6a).

**Figure 4.**
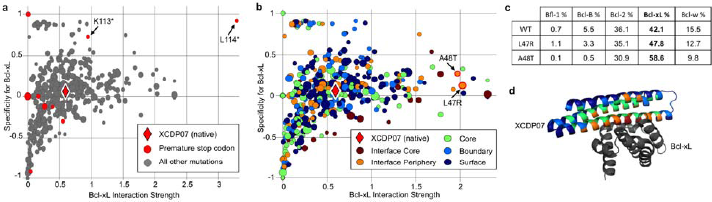
SynAg for large library protein interaction characterization. **(a,b)** Interaction strength versus specificity plots for a site-saturation mutagenesis library of the *de novo* Bcl-xL binder, XCDP07, with **(a)** premature stop codon variants in red and **(b)** two confirmed affinity and specificity improving mutations highlighted. For each protein variant, the diameter is a function of its representation in the naïve library, which is used as a measure of confidence, **(c)** The pro-survival BCL2 homologue mating distribution for wild-type (WT) XCDP07 and two point mutants that showed enrichment in a one-sided yeast surface display screen^21^, **(d)** A feature-colored cartoon model of XCDP07 bound to Bcl-xL.

Favorable mutations from a yeast surface display library were correctly identified using library-on-library SynAg, but with additional information about relative binding affinities and specificities (Fig. 4b,c). In particular, two mutations at the interface periphery, L47R and A48T, were found to improve interaction strength with Bcl-xL. Both mutations were enriched by fluorescence-activated cell sorting of an XCDP07 site-saturation mutagenesis surface display library incubated with fluorescently labeled Bcl-xL and unlabeled competitor homologues^21^. Unlike a traditional one-sided yeast surface display assay, SynAg provided detailed information about binding affinities and specificities to each target (Fig. 4c). We observed moderately improved on-target specificity for L47R, mostly through relative weakening of the interactions with Bcl-w and Bcl-B. We observed that A48T more dramatically weakened all off-target interactions with a 16.5% increase of on-target binding. SynAg was also able to confirm two favorable mutations in a Bcl-2 binder, 2CDP06, that were previously identified with yeast surface display enrichment^21^ (Fig. S6b).

### Characterizing the effect of environmental changes

To demonstrate the characterization of a protein interaction network in a new extracellular environment, we added a soluble competitive binder at the start of a batched mating, which selectively inhibited protein-protein interactions up to 800-fold (Fig. 5a,b). A pro-apoptotic BCL2 peptide, the BH3 domain of Bad (Bad.BH3), was found to interact selectively with five pro¬survival BCL2 homologues (Fig. 3f). No interaction was observed with Bcl-B. Library-on-library matings with and without 100 nM Bad.BH3 were normalized to one another with the assumption that interactions involving Bcl-B were unaffected. Normalization accounted for differences in total sequencing reads between conditions.

**Figure 5.**
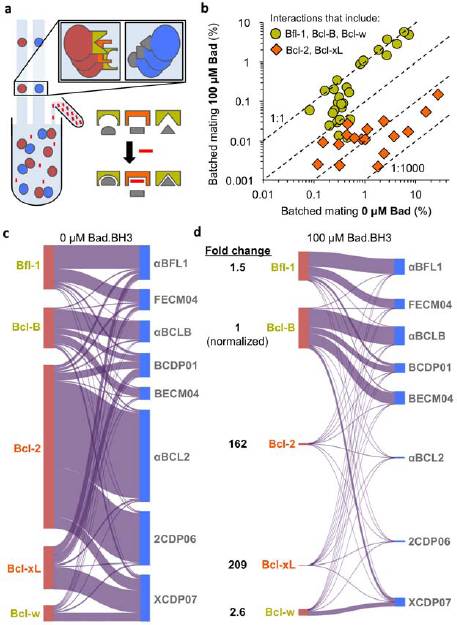
SynAg for characterizing the effect of environmental changes on protein interactions. **(a)** The agglutination environment can be manipulated with the addition of arbitrary molecules (red) that disrupt interactions involving particular proteins (orange) but not others (yellow), **(b)** The addition of a competing peptide, Bad.BH3, to a mating between five pro-survival BCL2 homologues and eight *de novo* binding proteins results in the isolated disruption of interactions involving the peptide’s binding targets, Bcl-2 and Bcl-xL (Fig. 3f). Dashed lines representing 1:1, 1:10, 1:100, and 1:1000 differences between conditions are given for reference. **(c,d)** The protein-protein interaction network visualized **(c)** without and **(d)** with the addition of 100 nM Bad.BH3. Fold changes are given for the aggregate interactions of each BCL2 homologue.

The addition of 100 μM Bad.BH3 resulted in specific inhibition of interactions involving its expected binding partners: Bcl-2 and Bcl-xL. Interactions involving these two homologues were inhibited by at least 16-fold and up to 800-fold (Fig. 5b,c,d). No change was observed for all strong protein-protein interactions involving pro-survival homologues that weakly interact with Bad.BH3: Bfl-1, Bcl-B, and Bcl-w. Weak protein-protein interactions involving these three homologues showed reduced mating efficiency in the presence of Bad.BH3, which can be attributed to an increased concentration of bulk protein in the media that serves to block non-specific interactions. Considered together, all protein-protein interactions involving Bcl-xL and Bcl-2 were strongly inhibited, with normalized mating percent fold changes of 209 and 162, respectively. The weaker Bad.BH3 binders, Bcl-w and Bfl-1, displayed a normalized mating percent fold change of 2.6 and 1.5, respectively. All aggregate fold changes were consistent with previous characterization of Bad.BH3 interactions with the five pro-survival homologues (Fig. 3f) and with previous work^19^.

## Discussion

We showed that the mating of *S. cerevisiae* can be reprogrammed by the surface expression of arbitrary SynAg proteins that replace the function of the native sexual agglutinin proteins, Aga2 and Sag1. Using SynAg, we demonstrated quantitative library-on-library characterization of up to 7000 distinct protein-protein interactions in a single pot. Additionally, we showed how SynAg could be used for characterizing protein interaction networks in different environments by adding an exogenous competitor to the mating environment. To date, tools for screening libraries of protein interactions are limited by throughput, a fixed intracellular environment, or accuracy. Previous strategies for developing library-on-library screening platforms have used cell-free systems, which are far less scalable. In contrast, SynAg combines the scalability of a cellular assay with the feature of environmental manipulation on a library-on-library scale.

The BCL2 protein network was chosen to demonstrate and validate SynAg due to previous characterization that showed a wide range of interaction strengths. We expect any of the diverse classes of proteins that can be functionally expressed on the surface of yeast^33^ to be compatible with SynAg. Some proteins do not functionally display on the yeast surface, and would therefore not be compatible with SynAg. For example, the detection of interactions requiring specific post-translational modifications may not be possible^34^. Additionally, SynAg is likely ill-suited for the screening of homodimer libraries. Oligomeric proteins are known to display on the yeast surface as functional assemblies,^35^ which means that homodimers would already be bound and not accessible for agglutination with a neighboring yeast cell expressing the same protein. Further studies are required to investigate these and other possible restrictions to SynAg.

SynAg provides a high-throughput platform for screening environment-responsive protein interactions and interaction-inhibiting drug candidates. Engineered protein-protein interactions that respond to environmental changes, such as pH, are valuable for biosensors^36^ and drug delivery^37^. SynAg may enable the rapid identification of functional variants using one-pot screening of design libraries rather than individual testing of protein pairs. SynAg also has potential applications for pharmaceutical development. Drug-induced protein interaction inhibition is a powerful therapeutic strategy for treating cancers, inflammation, and infectious diseases^38^. SynAg may streamline pre-clinical drug screening workflows by testing candidate compounds on protein interaction networks, enabling simultaneous screening for efficacy and specificity.

In addition to its utility for characterizing protein interactions, SynAg provides a unique ecological model for studying pre-zygotic genetic isolation. Previous work described the large diversity in sexual agglutination proteins across yeast species and suggested that co-evolution of these proteins may drive speciation by genetically isolating haploid pairs^39^. Here, we have created a fully engineerable synthetic pre-zygotic barrier that can be used as a model to study complex ecological phenomena such as speciation and sexual selection, similar to the use of engineered *E.coli*for modeling predator-prey dynamics^40^.

## Methods

### DNA Construction

Isogenic fragments for yeast transformation or plasmid assembly were PCR amplified from existing plasmids or yeast genomic DNA with Kapa polymerase (Kapa Biosystems), gel extracted from a plasmid digest (Qiagen), or synthesized by a commercial supplier (Integrated DNA Technologies). Plasmids were constructed with isothermal assembly^26^ and verified with Sanger sequencing^27^. Site-saturation mutagenesis library DNA was prepared with overlap PCR using custom NNK primers for each codon^21^. For a complete list of plasmids used in this study, see Table S7. Sequences for all cloning primers, fragments, and plasmids are available upon request.

### Yeast Methods

Unless otherwise noted, yeast transformations were performed with a standard lithium acetate transformation^41^ using approximately 300 ng of plasmid digested with PmeI. Yeast Peptone Dextrose (YPD), Yeast Peptone Galactose (YPG), and Synthetic Drop Out (SDO) medium supplemented with 80 mg/mL adenine were made according to standard protocols. Saturated yeast cultures were prepared by inoculating 3 mL of YPD from a freshly struck plate and growing for 24 hours at 30°C.

### Yeast Strain Construction

EBYlOOα was generated from a mating between EBYlOOa and a leucine prototroph W303α variant. Following sporulation and tetrad dissection, replica plating was used to identify MATα haploids auxotrophic for lys and trp^42^. Plating on 5-FOA was used to select strains with URA3 inactivating mutations^43^. Final ySYNAG strains were constructed with many rounds of chromosomal integration, each consisting of a single transformation, auxotrophic or antibiotic selection, and PCR to verify integration into the expected locus. For a complete list of yeast strains used in this study, see Table S8.

### Yeast Site-Saturation Mutagenesis Library Construction

Site-saturation mutagenesis libraries were transformed into yeast using nuclease assisted chromosomal integration (Fig. S4c). Prior to transformation, yeast strains were grown in YPG for five hours. Library transformations were conducted with 2μ of each fragment and 10x cells and reagent volumes. Cells were washed in 1 mL YPD and resuspended in YPD to a total volume of 5 mL. A dilution series was plated on SDO-trp to quantify transformation efficiency. The remaining culture was grown for 5 hours, washed twice with 5 mL SDO-trp, and grown in 20 mL SDO-trp for 17 hours. 2 mL 25% glycerol aliquots were stored at −80°C.

### Protein Purification

DNA encoding the BH3 domain of Bad (Bcl-2 agonist of cell death protein; residues 103-131) was inserted into a modified pMAL-c5x vector resulting in an N-terminal fusion to maltose binding protein and a C-terminal 6-histidine tag. The vector was transformed into BL21(DE3)* *E. coli* (NEB) for protein expression. Protein was purified from soluble lysate first with nickel affinity chromatography (NiNTA resin; Qiagen), then by size exclusion chromatography (Superdex 75 10/300 GL; GE). Purified protein was concentrated via centrifugal filter (Millipore), snap-frozen in liquid nitrogen and stored at −80°C.

### Surface Expression Screening

10μL of saturated cells were washed with 1 mL PBSF, incubated in 50μL PBSF media with 1μg FITC-anti-myc antibody (Immunology Consultants Laboratory, Inc.) for 1 hour at 22°C, washed with 1 mL PBSF, and read with the FL1.A channel on an Accuri C6 cytometer. SynAg expression cassettes contain an HA tag between Aga2 and the gene of interest that can be used as an alternative epitope tag for labeling SynAg variants containing premature stop codons.

### Pairwise Mating Assays

2.5 μL from a saturated MATa culture and 5 μL from a saturated MATα culture were combined in 3 mL of YPD media and incubated at 30°C and 275 RPM for 17 hours. 5 μL were added to 1 mL of water and cellular expression of mCherry and mTurquoise was characterized with a Miltenyi MACSQuant VYB cytometer using channels Y2 and V1, respectively. A standard yeast gate was applied to all cytometry data and Flowjo was used for analysis and visualization.

### Yeast Library Preparation

Pre-characterized yeast libraries were prepared by manually combining individually transformed isogenic yeast strains with known barcodes (Table S1). Saturated isogenic yeast strains of the same mating type were pooled with equal cell counts, measured with an Accuri c6 flow cytometer.

Next-generation sequencing was used to characterize the barcodes and relative population distribution of yeast site-saturation mutagenesis libraries. A 2 mL glycerol stock was thawed, washed once with 1 mL YPD, and grown in 50 mL YPD for 24 hours. Genomic DNA was then prepared for next-generation sequencing.

### Library-on-Library Mating Assays

2.5 μL of saturated MATa library and 5 μL of saturated MATα library were combined in 3 mL of YPD and incubated at 30°C and 275 RPM for 17 hours. When characterizing interactions in the presence of Bad.BH3, the peptide was added at a concentration of 100 nM. 1 mL was washed twice in 1 mL SDO-lys-leu and transferred to 50 mL SDO-lys-leu with 100 nM β-estradiol for 24 hours. Genomic DNA was prepared for next-generation sequencing.

### Preparation for Next-Generation Sequencing

50 mL yeast cultures were harvested by centrifugation and lysed by heating to 70°C for 10 min in 2 mL 200 mM LiOAc and 1% SDS^44^. Cellular debris was removed with centrifugation and the supernatant was incubated at 37°C for 4 hours with 0.05 mg/mL RNase A. Following an ethanol precipitation, a 2% agarose gel was run to verify genomic DNA extraction. Two rounds of qPCR were performed to amplify a fragment pool from the genomic DNA and to add standard Illumina sequencing adaptors and assay specific index barcodes. Both PCRs were terminated before saturation in order to minimize PCR bias. The first PCR was run for 25-30 cycles, and the second PCR was run for 5-7 cycles. The final amplified fragment was gel extracted, quantified with a Qubit and sequenced with a MiSeq sequencer (Illumina).

### Accession numbers

BioProject Accession number: PRJNA380247

BioSample Accession numbers: SAMN06642476, SAMN06642477, SAMN06642478, SAMN06642479, SAMN06642480, SAMN06642481, SAMN06642482, SAMN06642483, SAMN06642484, SAMN06642485

### Code availability

All code for sequencing analysis is fully available on GitHub: https://github.com/dyounger/yeast_synthetic_agglutination

Sankey diagrams were generated with sankeyMATIC (http://sankeymatic.com/)

## Acknowledgements

We thank M. Dunham for technical discussions, A. Rosenberg for data analysis support and M. Parks and the UW Biofab for assistance with the construction of many plasmids and yeast strains used in this study. This work was supported by US National Science Foundation (NSF) award number 1317653. D.Y. and S.B. are supported by the NSF GRFP.

## Author Information

### Contributions

D.Y., D.B., and E.K. contributed to the technical design. D.Y. and S.B. implemented the methods. D.Y. analyzed the data and wrote the manuscript with contributions from all authors.

## Supplementary Information

**Figure S2.**
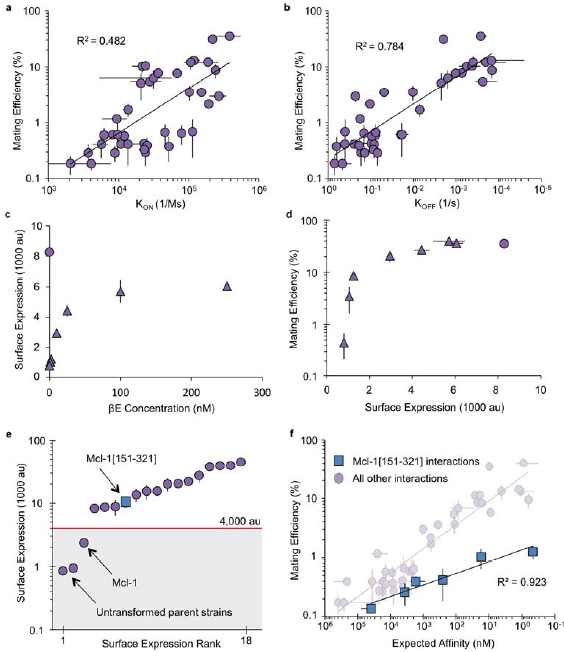
Pairwise SynAg mating efficiencies are affinity dependent. **(a,b)**Binding dynamics were previously measured with biolayer interferometry^21^. Here, we showed that both **(a)** on-rate and **(b)** off-rate both correlated with pairwise SynAg mating efficiency (SD, n=3). **(c,d,e,f)** We analyzed the effect of variation in surface expression strength, **(c)** A β-Estradiol (βE) inducible SynAg expressing strain was characterized at a range of inducer concentrations (triangles) along with a constitutive SynAg expressing strain without induction (circle) (SD, n=2). **(d)** SynAg assays were conducted to compare mating efficiency with surface expression strength. Above 4,000 au, minimal changes in mating efficiency were observed (SD, n=2). **(e)** Of the BCL2 homologues and *de novo* binding proteins, a single SynAg protein, Mcl-1, had a surface display strength below 4,000 au. A semi-functional alternative truncation improved surface display strength (blue square) (SD, n=2). **(f)** A SynAg assay was used to characterize the interaction strength between the semi-functional truncation and *de novo* binding proteins. With improved surface expression strength, mating recovery was observed. However, the truncation seemed to decrease the affinity for all binders (SD, n=3).

**Figure S3.**
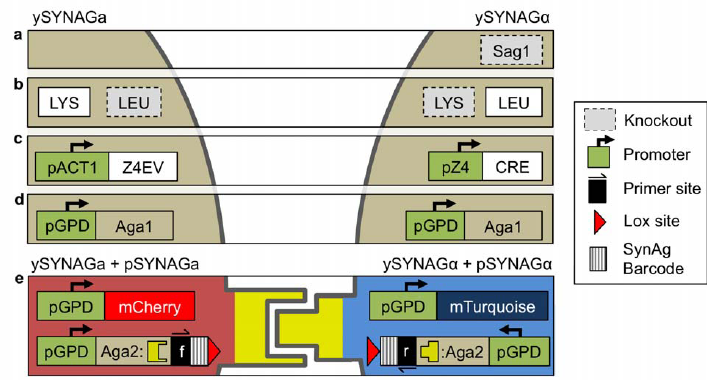
Yeast strain genetic components. **(a)** The α-agglutinin, Sag1, is knocked out in ySYNAGato eliminate wild-type agglutination, **(b)** ySYNAGa and ySYNAGα have complementary lysine and leucine markers for diploid selection, **(c)** ySYNAGa cells constitutively express a (β-Estradiol inducible transcription factor, Z4EV, which activates the pZ4 promoter for CRE recombinase expression in diploid cells^22^, **(d)** ySYNAGa and ySYNAGα constitutively express Agal for yeast surface display. The strongest *S. cerevisiae*-native constitutive promoter, pGPD, was chosen to maximize expression, **(e)** A transformation with pSYNAGa or pSYNAGα adds a constitutively expressed fluorescent reporter and SynAg protein fused to Aga2. A recombination site, barcode, and primer binding site flank the SynAg expression cassettes.

**Figure S4.**
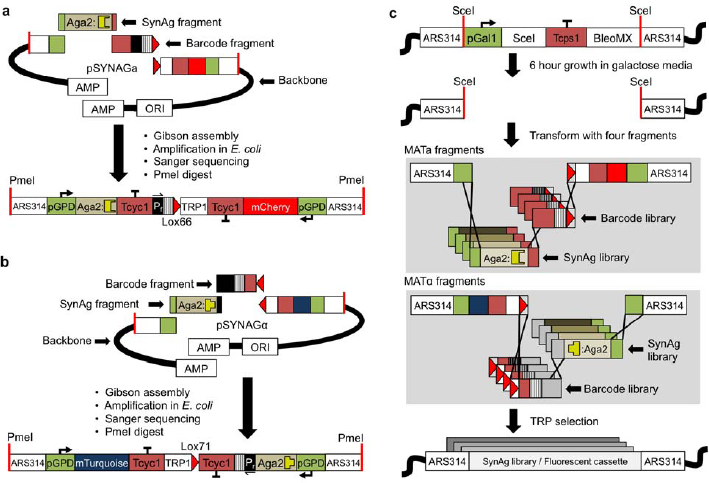
Construction strategy for SynAg plasmids and yeast strains. **(a)** pSYNAGa and **(b)** pSYNAGα were each constructed with a four-fragment Gibson assembly. Fragments included a SynAg fragment, a barcode-containing fragment amplified with a degenerate primer, and two backbone fragments. The strongest *S. cerevisiae*-native constitutive promoter, pGPD, was chosen to maximize expression of both the SynAg expression cassette and fluorescent reporter. Following transformation into *E. coli,* plasmid open reading frames and barcodes were Sanger sequenced. Verified plasmids were digested with a restriction enzyme, Pmel, and transformed into ySYNAGa or ySYNAGα. **(c)** For library integrations, ySYNAGa and ySYNAGα were grown for hours in galactose media to induce expression of an endonuclease, Seel, which caused DNA damage at the integration site. Cells were then transformed with four mating type specific fragments that recombine and integrate into the chromosome.

**Figure S5.**
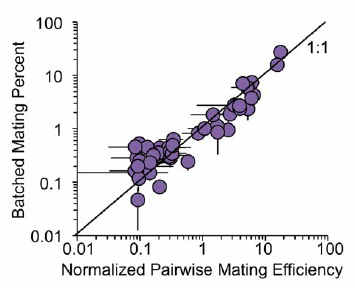
A comparison of pairwise and library-on-library SynAg assays. Mating efficiency from a pairwise SynAg assay was normalized such that the sum of all mating efficiencies was 100. A line showing a 1:1 relationship is given for reference. Error bars represent one standard deviation from at least two replicates. Bcl-2 Interaction Strength

**Figure S6.**
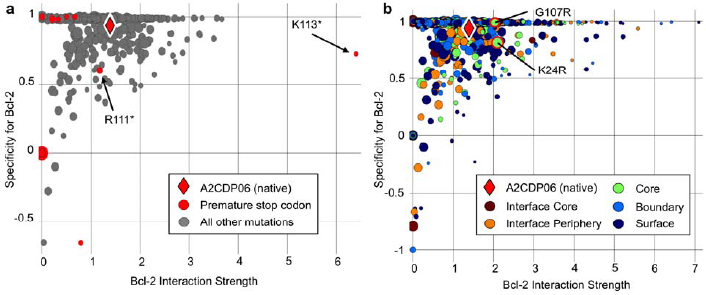
Site-saturation mutagenesis library analysis of a *de novo* Bcl-2 binder. Interaction strength versus specificity plots for a site-saturation mutagenesis library of the 116-residue 2CDP06^21^ with **(a)** premature stop codon variants in red and **(b)** two confirmed affinity-improving mutations highlighted. For each protein variant, the diameter is a function of its representation in the naïve library, which is used as a measure of confidence.

**Table S1.**
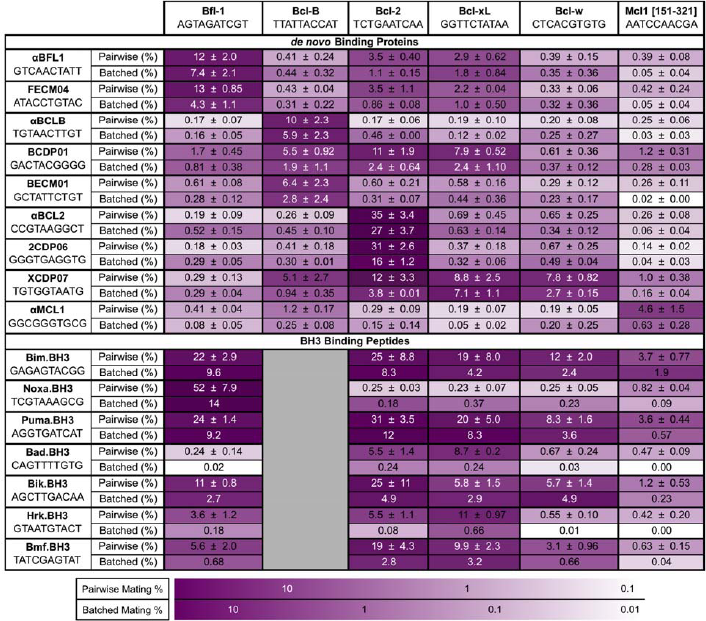
Characterization of protein interactions using pairwise and batched synthetic agglutination. Interactions involving six pro-survival BCL2 homologues (columns), nine *de novo* binding proteins (upper rows), and seven BH3 binding peptides (lower rows) were characterized with pairwise and batched SynAg assays. The unique 10 bp barcode used to identify each SynAg protein with next-generation sequencing is also listed. For each interaction, the pairwise mating percent is given with an error of one standard deviation (n=3). The batched mating percent for interactions involving the *de novo* binding proteins are given with an error of one standard deviation (n=2). Shading provides a qualitative comparison between the pairwise and batched SynAg methods.

**Table S7.**
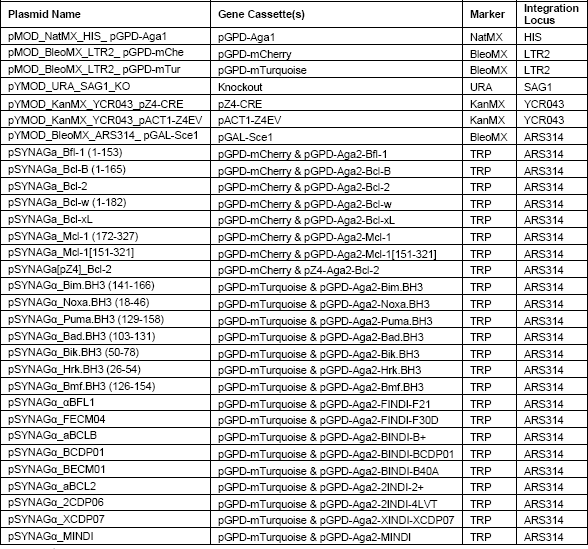
Plasmids used in this study. Common truncation for some pro-survival BCL2 homologues and pro-apoptotic peptides were used and are noted in parenthesis with the plasmid name where applicable.

**Table S8.**
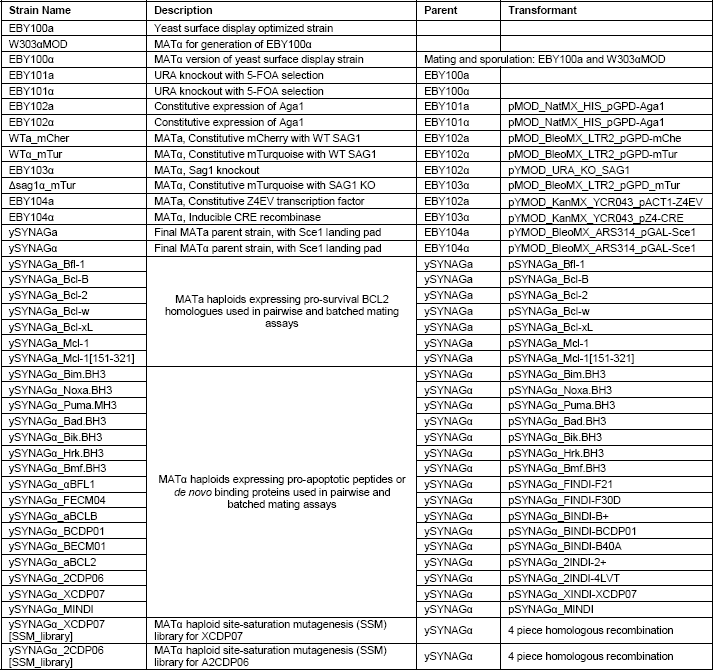
Yeast strains and yeast libraries used in this study.

## References

1. Fields, Stanley, and Ok-kyu Song (1989) A novel genetic system to detect protein protein interactions. Nature 340.6230: 245–246.

2. Yu, Haiyuan, et al. (2011) Next-generation sequencing to generate interactome datasets. Nature Methods 8.6: 478–480.

3. Hastie, Alex R., and Steven C. Pruitt (2007) Yeast two-hybrid interaction partner screening through in vivo Cre-mediated Binary Interaction Tag generation. Nucleic Acids Research 35.21: e141.

4. Yachie, Nozomu, et al. (2016) Pooled-matrix protein interaction screens using Barcode Fusion Genetics. Molecular Systems Biology 12.4: 863.

5. Chen, Yu-Chi, et al. (2010) Exhaustive benchmarking of the yeast two-hybrid system. Nature methods 7.9: 667–668.

6. Braun, Pascal, et al. (2009) An experimentally derived confidence score for binary protein-protein interactions. Nature Methods 6.1: 91–97.

7. Smith, George (1985) Filamentous fusion phage: novel expression vectors that display cloned antigens on the virion surface. Science 228: 1315–1318.

8. Boder, Eric, et al. (1997) Yeast surface display for screening combinatorial polypeptide libraries. Nature Biotechnology 15.6: 553–557.

9. Perfetto, Stephen P., Pratip K. Chattopadhyay, and Mario Roederer (2004) Seventeen-colour flow cytometry: unravelling the immune system. Nature Reviews Immunology 4.8: 648–655.

10. Gu, Liangcai, et al. (2014) Multiplex single-molecule interaction profiling of DNA-barcoded proteins. Nature 515.7528: 554–557.

11. Roy, A. M. I. T., et al. (1991) The AGA1 product is involved in cell surface attachment of the Saccharomyces cerevisiae cell adhesion glycoprotein a-agglutinin. Molecular and Cellular Biology 11.8: 4196–4206.

12. Dranginis, A et al. (2007) A Biochemical Guide to Yeast Adhesins: Glycoproteins for Social and Antisocial Occasions. Microbiology and Molecular Biology Reviews 71.2: 282–294.

13. Zhao, Hui, et al. (2001) Interaction of α-agglutinin and a-agglutinin, Saccharomyces cerevisiae sexual cell adhesion molecules. Journal of Bacteriology 183.9: 2874–2880.

14. Herskowitz, Ira. (1988) Life cycle of the budding yeast Saccharomyces cerevisiae. Microbiological Reviews 52.4: 536–553.

15. Boder, Eric, Katarina Midelfort, and Dane Wittrup. (2000) Directed evolution of antibody fragments with monovalent femtomolar antigen-binding affinity. Proc Natl Acad Sci USA 97.20: 10701–10705.

16. Huang, Dagang, and Eric Shusta. (2005) Secretion and surface display of green fluorescent protein using the yeast Saccharomyces cerevisiae. Biotechnology Progress 21.2: 349–357.

17. Kim, Yong-Sung, et al. (2006) Directed evolution of the epidermal growth factor receptor extracellular domain for expression in yeast. PROTEINS: Structure, Function, and Bioinformatics 62.4: 1026–1035.

18. Adams, Jerry M., and Suzanne Cory. (1998) The Bcl-2 protein family: arbiters of cell survival. Science 281.5381: 1322–1326.

19. Chen, Lin, et al. (2005) Differential targeting of prosurvival Bcl-2 proteins by their BH3-only ligands allows complementary apoptotic function. Molecular Cell 17.3: 393–403.

20. Procko, Erik, et al. (2014) A computationally designed inhibitor of an Epstein-Barr viral Bcl-2 protein induces apoptosis in infected cells. Cell 157.7: 1644–1656.

21. Berger, Stephanie, et al. (2016) Computationally designed high specificity inhibitors delineate the roles of BCL2 family proteins in cancer. eLife 5: e20352.

22. Mclsaac, R. Scott, et al. (2013) Synthetic gene expression perturbation systems with rapid, tunable, single-gene specificity in yeast. Nucleic Acids Research: 41.4: e57.

23. Day, Catherine L., et al. (2005) Solution structure of prosurvival Mcl-1 and characterization of its binding by proapoptotic BH3-only ligands. Journal of Biological Chemistry 280.6: 4738–4744.

24. Sauer, Brian, and Nancy Henderson (1988) Site-specific DNA recombination in mammalian cells by the Cre recombinase of bacteriophage P1. Proc Natl Acad Sci USA 85.14: 5166–5170.

25. Louvion, Jean-François, Biserka Havaux-Copf, and Didier Picard (1993) Fusion of GAL4-VP16 to a steroid-binding domain provides a tool for gratuitous induction of galactose-responsive genes in yeast. Gene 131.1: 129–134.

26. Gibson, Daniel G., et al. (2009) Enzymatic assembly of DNA molecules up to several hundred kilobases. Nature Methods 6.5: 343–345.

27. Sanger, Frederick, Steven Nicklen, and Alan R. Coulson (1977) DNA sequencing with chain-terminating inhibitors. Proc Natl Acad Sci USA 74.12: 5463–5467.

28. Araki, Kimi, Masatake Araki, and Ken-ichi Yamamura. (2002) Site-directed integration of the cre gene mediated by Cre recombinase using a combination of mutant lox sites. Nucleic Acids Research 30.19: e103.

29. Bentley, David R., et al. (2008) Accurate whole human genome sequencing using reversible terminator chemistry. Nature 456.7218: 53–59.

30. Van Delft, Mark F., and David CS Huang (2006) How the Bcl-2 family of proteins interact to regulate apoptosis. Cell Research 16.2: 203–213.

31. Lee, Michael E., et al. (2015) A highly characterized yeast toolkit for modular, multipart assembly. ACS Synthetic Biology 4.9: 975–986.

32. Wingler, Laura M., and Virginia W. Cornish (2011) Reiterative recombination for the in vivo assembly of libraries of multigene pathways. Proc Natl Acad Sci USA 108.37: 15135–15140.

33. Gai, S. Annie, and K. Dane Wittrup (2007) Yeast surface display for protein engineering and characterization. Current Opinion in Structural Biology 17.4: 467–473.

34. Wadle, Andreas, et al. (2005) Serological identification of breast cancer-related antigens from a Saccharomyces cerevisiae surface display library. International Journal of Cancer 117.1: 104–113.

35. van den Beucken, Twan, et al. (2003) Affinity maturation of Fab antibody fragments by fluorescent-activated cell sorting of yeast-displayed libraries. FEBS Letters 546.2-3: 288–294.

36. Strauch, Eva-Maria, Sarel J. Fleishman, and David Baker (2014) Computational design of a pH-sensitive IgG binding protein. Proc Natl Acad Sci USA 111.2: 675–680.

37. Lee, Eun Jung, Na Kyeong Lee, and In-San Kim (2016) Bioengineered protein-based nanocage for drug delivery. Advanced Drug Delivery Reviews 106: 157–171.

38. Wells, James A., and Christopher L. McClendon (2007) Reaching for high-hanging fruit in drug discovery at protein-protein interfaces. Nature 450.7172: 1001–1009.

39. Xie, Xianfa, et al. (2011) Accelerated and adaptive evolution of yeast sexual adhesins. Molecular Biology and Evolution 28.11: 3127–3137.

40. Balagaddé, Frederick K., et al. (2008) A synthetic Escherichia coli predator-prey ecosystem. Molecular Systems Biology 4.1: 187.

41. Gietz, R.Daniel, and Robert H. Schiestl (2007) High-efficiency yeast transformation using the LiAc/SS carrier DNA/PEG method. Nature Protocols 2.1: 31–34.

42. Spencer, John FT, Dorothy M. Spencer, and I. J. Bruce (2012) Yeast Genetics: A Manual of Methods. Springer Science & Business Media.

43. Boeke, Jef D., Francois Croute, and Gerald R. Fink (1984) A positive selection for mutants lacking orotidine-5-phosphate decarboxylase activity in yeast: 5-fluoro-orotic acid resistance. Molecular and General Genetics 197.2: 345–346.

44. Lõke, Marko, Kersti Kristjuhan, and Arnold Kristjuhan (2011) Extraction of genomic DNA from yeasts for PCR-based applications. Biotechniques 50.5: 325–328.

